# Extreme diversity of mycoviruses present in four strains of *Rhizoctonia cerealis*, the pathogen of wheat sharp eyespot

**DOI:** 10.1101/2022.11.01.514804

**Authors:** Wei Li, Haiyan Sun, Shulin Cao, Aixiang Zhang, Haotian Zhang, Yan Shu, Huaigu Chen

## Abstract

*Rhizoctonia cerealis* is the pathogen of wheat sharp eyespot, which occurs throughout temperate wheat growing regions of the world. In this project, the genomes of viruses from four strains of *R. cerealis* were analyzed based on Illumina high-throughput RNA-Seq data. Ribosomal RNA-depleted total RNA and purified dsRNA from cultivated mycelia of each isolate were used for cDNA library construction and sequencing. After filtering out reads that mapped to the fungal genome, viral genomes were assembled using the remaining reads from the rRNA-depleted and dsRNA-Seq data. In total, 131 viral genome sequences containing complete ORFs, belonging to 117 viruses, were obtained. Based on phylogenetic analysis, some of them were identified as novel members of the families *Curvulaviridae, Endornaviridae, Hypoviridae, Mitoviridae, Mymonaviridae* and *Phenuiviridae*, while others were unclassified viruses. We compared the integrity and reliability of the viral sequences obtained by the two sequencing methods and, for the first time, estimated the density of some viruses in host cells. Most of these viruses from *R. cerealis* were sufficiently different from those deposited in databases. We propose the establishment of a new family, *Rhizoctobunyaviridae*, and two new genera, *Rhizoctobunyavirus* and *Iotahypovirus*. We further clarified the distribution and co-infection of these viruses in the four *R. cerealis* strains. In conclusion, the diversity of mycoviruses in *R. cerealis* is extremely rich. We report a series of novel viruses and provide important insight into virus evolution.

## Introduction

Viruses are the most abundant and diverse species on Earth and exist in almost all known cellular life [1]. With the rapid development of high-throughput sequencing and bioinformatics analysis technology, meta-virome studies could identify many viral genomes from a specific environment or biological population samples, and a large number of new viruses have recently been discovered from soil, water, ocean, intestine, invertebrate and mammalian samples [2–4]. Mycoviruses are common in major filamentous fungi, and some of them can reduce the pathogenicity of their host fungi, providing new resources for the control of fungal diseases in plants and animals [5–8].

Recent research has shown that the DNA mycovirus Sclerotinia sclerotiorum hypovirulence-associated DNA virus 1 (SsHADV-1) can convert its host, *S. sclerotiorum*, from a typical necrotrophic pathogen to a beneficial endophytic fungus [9]. It is not unique in this ability; the dsRNA mycovirus Pestalotiopsis theae Chrysovirus 1 (PtCV1) can also convert pathogenic *P. theae* to an endophytic fungus, and the presence of PtCV1 confers host plants with high resistance against virulent *P. theae* strains [10]. The effects of mycoviruses on controlling the endophytic traits of host fungi suggest that viruses may play a special role in ecosystems composed of mycoviruses, pathogenic fungi and plants. These studies showed broad prospects for the exploitation and utilization of mycovirus resources.

*Rhizoctonia* is a form genus in the family *Ceratobasidiaceae*, which contains two genera, *Ceratobasidium* and *Thanatephorus* [11–13]. The diversity of mycoviruses in *R. solani* (teleomorph *Thanatephorus cucumeris*) has been widely reported [14–18]. *R. cerealis* (teleomorph *Ceratobasidium cereale*), the pathogen of wheat sharp eyespot, is widely distributed in temperate wheat-growing regions worldwide and causes major wheat yield losses [19]. We previously reported a mitovirus named Rhizoctonia cerealis mitovirus (RcMV1) and 22 endornaviruses from *R. cerealis* [20–22]. The results of dsRNA gel electrophoresis indicated the prevalence of mycoviruses in some *R. cerealis* strains [23].

Recently, we obtained a chromosome-level draft genome of *R. cerealis* [24], facilitating omics research on this fungus and the mycoviruses it harbors. In this study, the genomes of mycoviruses in four strains of *R. cerealis* were analyzed based on high-throughput RNA-Seq data. This project aims to understand the diversity, evolution and spread of mycoviruses in different strains of *R. cerealis*.

## Materials and Methods

### Fungal strains, total RNA and dsRNA extraction and Illumina sequencing

Four *R. cerealis* strains, R0928, R0942, R1084 and R10125, were collected from Henan, Anhui and Shandong Provinces in China [22]. Total RNA was extracted from fungal mycelium using the RNAprep Pure Plant Plus Kit (Tiangen, China). Ribosomal RNA was depleted using the Ribo-Zero™ Kit (Epicentre, Madison, WI, USA). The viral dsRNA of these strains was extracted and prepared as previously described [22]. Library preparation and Illumina sequencing were performed by Genepioneer Bio-Tech Co., Ltd., (China) in rRNA-depleted samples and Shanghai Hanyu Bio-Tech Co., Ltd., (China) in dsRNA samples. Each strain was subjected to rRNA-deletion RNA-Seq and dsRNA-Seq.

### Genome sequence assembly and confirmation

Before viral genome assembly, we filtered the reads from the host fungal genome. Reads from the clean RNA-Seq data were mapped to the genome of *R. cerealis* strain R0301 [24] with Bowtie2 [25] and SAMtools [26], and unmatched reads were extracted and assembled de novo using SOAPdenovo2 v2.04 [27] and Geneious Prime 2022.1.1 [28]. The contigs of viral origin were selected based on BLAST searches against the Nr database of the National Center for Biotechnology Information (NCBI). Viral contigs obtained from rRNA-deletion RNA-Seq and dsRNA-Seq were combined into one total viral sequence database. Each viral contig was used as a query for a BLAST search against the total viral sequence database to identify contigs overlapping the extended region. For each virus of each strain, all identified contigs were assembled again using the Geneious program. After several iterations, the viral genomes were finally determined.

The number of reads covering the viral genomes was obtained by mapping the reads from each sequenced library on reference sequences with Bowtie2 [25] and SAMtools [26]. Mapping results were displayed using the Geneious program. Based on these results, the reads per kilobase per million mapped reads (RPKM) value and depth of coverage (= (bases of all mapped reads)/(length of virus)) of each virus were calculated. For viruses with a depth of coverage less than 20, we designed specific primers, amplified the genome sequences by RT–PCR and sequenced them to confirm the viral genomes.

### Genome structure and phylogeny analysis

The open reading frames (ORFs) in the viral genomes were predicted using the Geneious program. The conserved domains and similar sequences of the deduced amino acid sequences were searched using BLASTp in the NCBI Conserved Domain Database, and domain hits with expected values less than 0.01 were considered (http://www.ncbi.nlm.nih.gov/Structure/cdd/wrpsb.cgi, e-value < 0.01). The genome structure of the viruses was displayed using the Geneious program.

The genome sequences of the viruses used in phylogenetic analysis were obtained from GenBank. Some of them were selected based on BLAST searches using the novel viruses in this study as a query. Other viruses representing different taxa were selected from the 2021 ICTV taxonomy release (#37) available online at https://ictv.global/taxonomy. Multiple alignments of the amino acid sequences of the viral RNA-dependent RNA polymerase (RdRp) regions were performed using the MAFFT program (https://myhits.isb-sib.ch/cgi-bin/mafft) [29] in the Geneious program. ModelFinder [30] was used to select the best-fit model by using the Bayesian information criterion. Maximum likelihood phylogenies were generated via ultrafast bootstrap analysis in which 1000 replicates were inferred using IQ-TREE [31], performed in the PhyloSuite v1.2.2 program [32]. The display, annotation, and management of phylogenetic trees were performed using the online Interactive Tree of Life tool (https://itol.embl.de/).

## Results

### Genome sequence assembly and confirmation

After several iterations of assembly, a total of 131 complete or almost complete viral genome sequences of 117 viruses were identified in the four *R. cerealis* strains (Table 1). Most of the viral genomes had only one segment, while 14 orthocurvulaviruses had two segments. Among these viruses, RcMV1 in R1084 and 20 endornaviruses in R0928, R0942, R1084 and R10125 have been reported previously [21, 22]. All of the new genome sequences of the viruses have been deposited in the NucBank in the National Genomics Data Center, Beijing Institute of Genomics, Chinese Academy of Sciences/China National Center for Bioinformation, under accession numbers C_AA001139 to C_AA001248, which are publicly accessible at https://ngdc.cncb.ac.cn/nucbank.

**Table 1.**
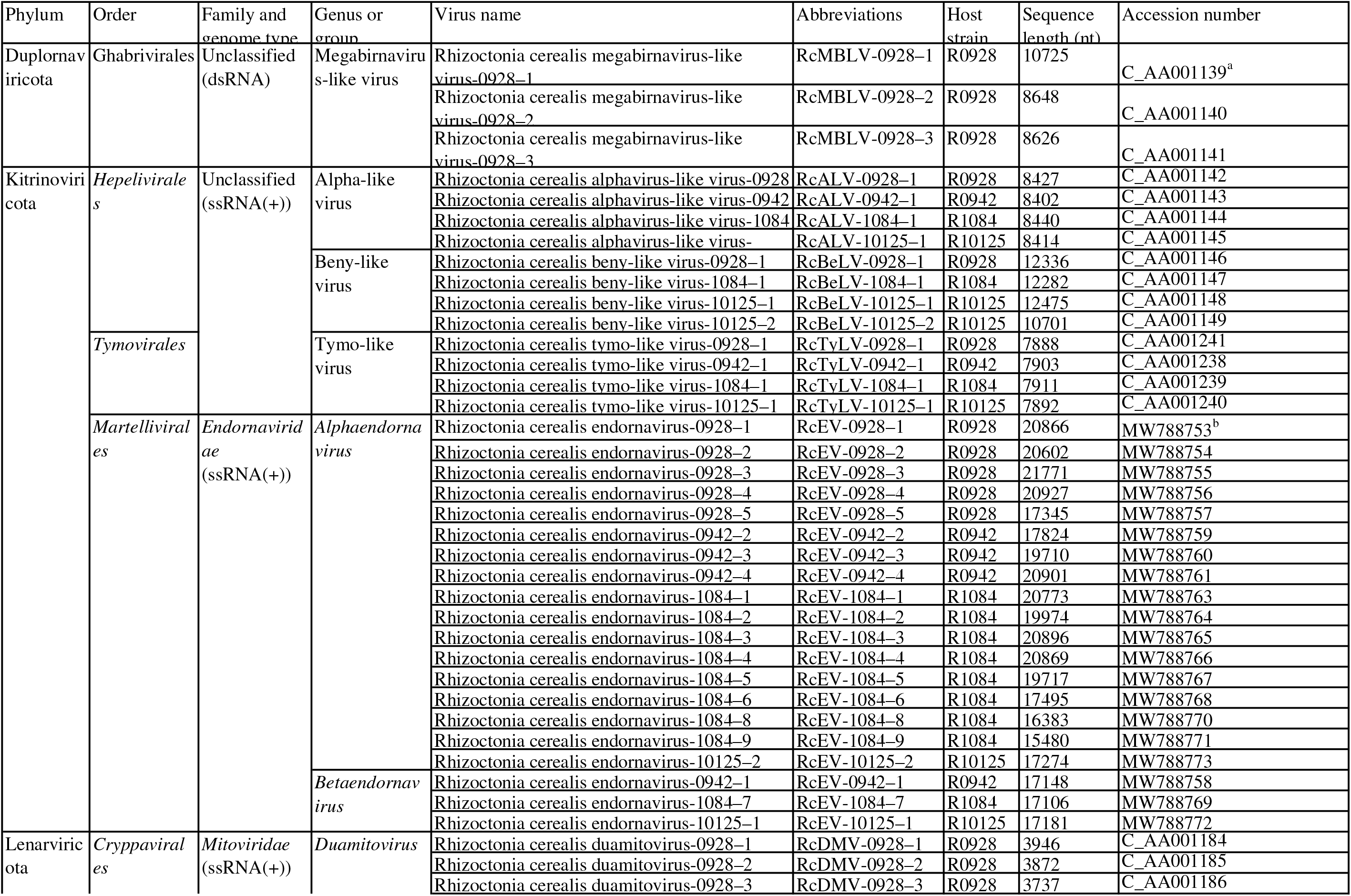
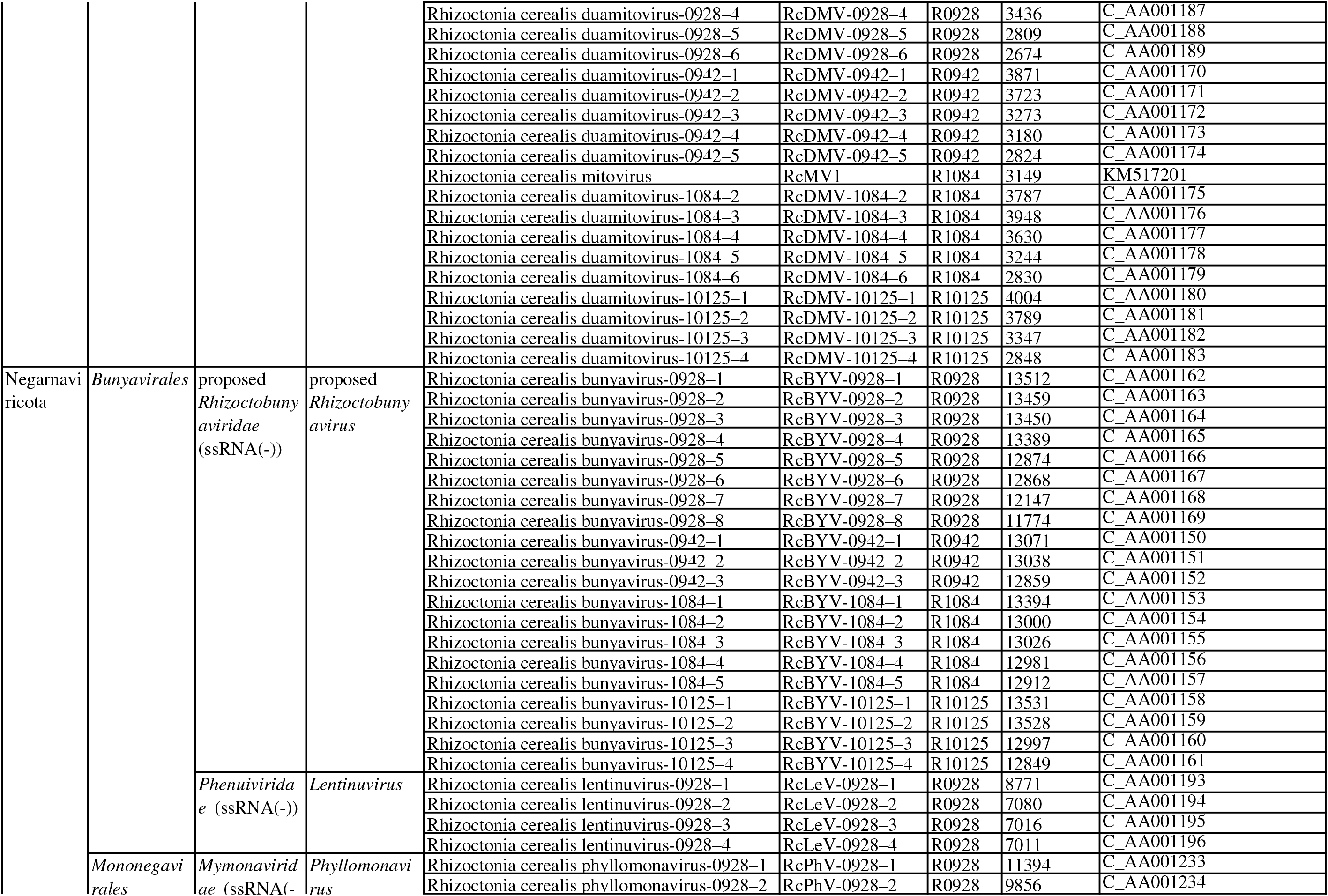
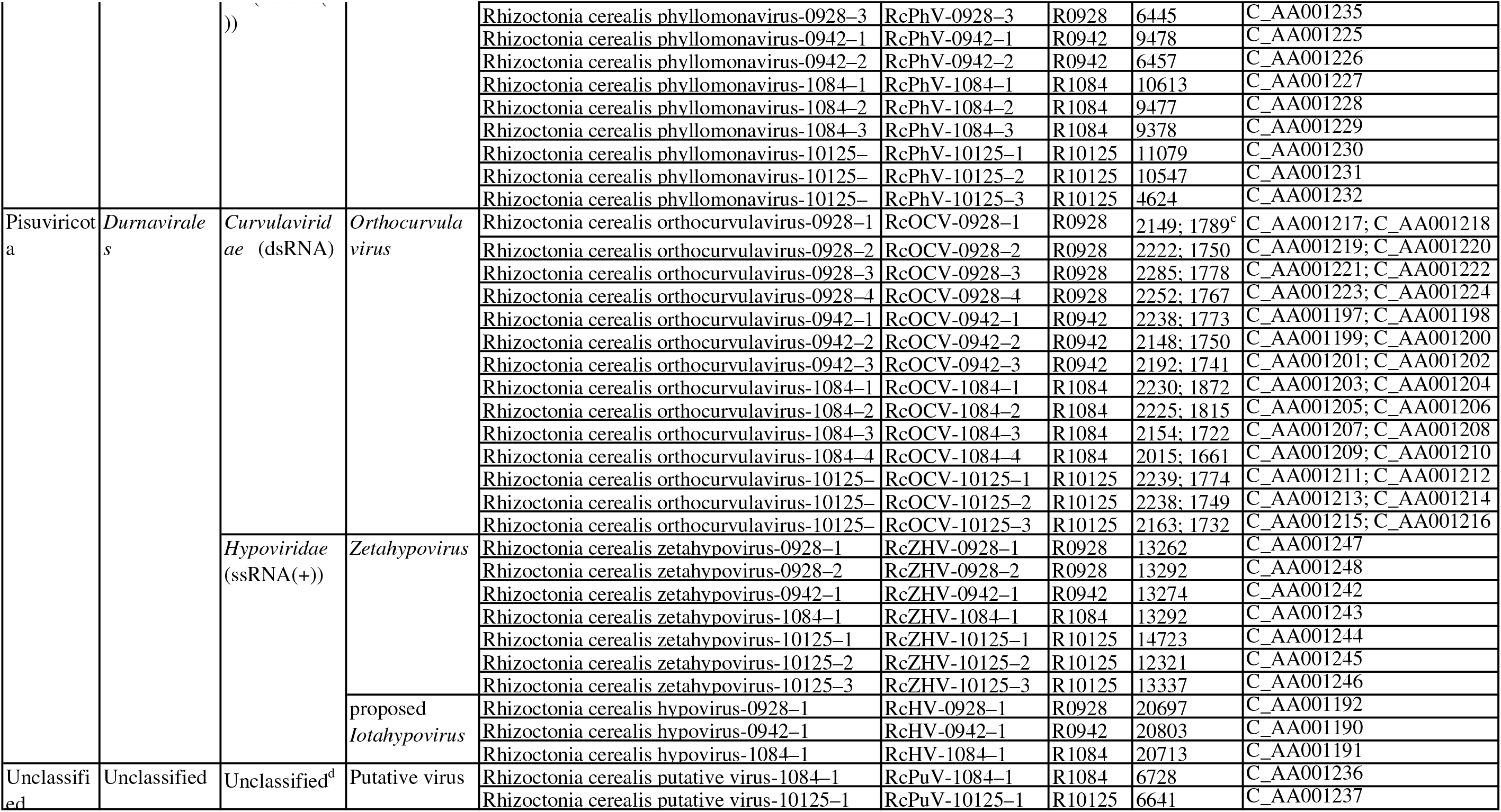
Information on viruses identified in the four *R. cerealis* strains. ^a^The accession number in NucBank. bThe accession number in GenBank. ^c^The length of RNA1 and RNA2 of the orthocurvulavirus. ^d^The genome type of this putative virus is unknown.

We calculated the RPKM value of each viral genome sequence based on the rRNA-depleted RNA-Seq and dsRNA-Seq data. For most viral genome sequences, the RPKM values obtained based on dsRNA-Seq data were higher than those based on rRNA-depleted RNA-Seq data (Table 2). However, for mitoviruses, higher RPKM values were obtained based on RNA-depleted RNA-seq data (Table 2). After combining these two datasets, we calculated the depth of coverage of each viral genome sequence. Most viral genomes showed high coverage, and only 13 of them showed less than 100-fold coverage (Table 2). Seven sequences with less than 20-fold coverage were further verified by RT–PCR and Sanger sequencing to confirm that all viral genome sequences were reliable.

**Table 2.** Number of reads mapped to viral genomes, RPKM, and depth of coverage for each virus based on dsRNA-Seq and rRNA-depleted RNA-Seq data.

| Viruses | Sequence length | dsRN-Seq |  | rRNA-depleted RNA- |  | Depth of coverage |
| --- | --- | --- | --- | --- | --- | --- |
|  |  | No. of reads | RPKM | No. of reads | RPKM |  |
| RcMBLV-0928-1 | 10725 | 31554 | 175.80 | 631 | 1.31 | 450.14 |
| RcMBLV-0928-2 | 8648 | 94104 | 650.20 | 38 | 0.10 | 1632.90 |
| RcMBLV-0928-3 | 8626 | 9301 | 64.43 | 0 | 0.00 | 161.74 |
| RcALV-0928-1 | 8427 | 13225 | 93.77 | 12 | 0.03 | 235.62 |
| RcALV-0942-1 | 8402 | 15776 | 114.19 | 271 | 0.79 | 286.49 |
| RcALV-1084-1 | 8440 | 38921 | 225.48 | 0 | 0.00 | 691.72 |
| RcALV-10125-1 | 8414 | 1644 | 11.01 | 16 | 0.04 | 29.59 |
| RcBeLV-0928-1 | 12336 | 21065 | 102.03 | 4701 | 8.49 | 313.30 |
| RcBeLV-1084-1 | 12282 | 24951 | 99.33 | 15656 | 21.68 | 495.93 |
| RcBeLV-10125-1 | 12475 | 19673 | 88.83 | 13641 | 24.87 | 400.57 |
| RcBeLV-10125-2 | 10701 | 13766 | 72.46 | 13049 | 27.74 | 375.88 |
| RcTyLV-0928-1 | 7888 | 5698 | 43.16 | 702 | 1.98 | 121.70 |
| RcTyLV-0942-1 | 7903 | 1915 | 14.74 | 878 | 2.71 | 53.01 |
| RcTyLV-1084-1 | 7911 | 4232 | 26.16 | 912 | 1.96 | 97.54 |
| RcTyLV-10125-1 | 7892 | 12398 | 88.49 | 1612 | 4.65 | 266.28 |
| RcEV-0928-1 | 20866 | 244152 | 699.15 | 155 | 0.17 | 1756.26 |
| RcEV-0928-2 | 20602 | 558927 | 1621.06 | 207 | 0.22 | 4070.97 |
| RcEV-0928-3 | 21771 | 951790 | 2612.25 | 218 | 0.22 | 6559.24 |
| RcEV-0928-4 | 20927 | 21583 | 61.63 | 4 | 0.00 | 154.73 |
| RcEV-0928-5 | 17345 | 54149 | 186.54 | 20 | 0.03 | 468.45 |
| RcEV-0942-1 | 17148 | 656865 | 2329.62 | 26793 | 38.13 | 5980.21 |
| RcEV-0942-2 | 17824 | 223924 | 764.04 | 611 | 0.84 | 1889.60 |
| RcEV-0942-3 | 19710 | 231338 | 713.81 | 658 | 0.81 | 1765.57 |
| RcEV-0942-4 | 20901 | 132591 | 385.81 | 68 | 0.08 | 952.05 |
| RcEV-1084-1 | 20773 | 189500 | 446.05 | 0 | 0.00 | 1368.36 |
| RcEV-1084-2 | 19974 | 571647 | 1399.38 | 399 | 0.34 | 4295.93 |
| RcEV-1084-3 | 20896 | 32347 | 75.69 | 0 | 0.00 | 232.20 |
| RcEV-1084-4 | 20869 | 102959 | 241.23 | 0 | 0.00 | 740.04 |
| RcEV-1084-5 | 19717 | 752391 | 1865.85 | 979 | 0.84 | 5731.37 |
| RcEV-1084-6 | 17495 | 2148281 | 6004.14 | 0 | 0.00 | 18419.10 |
| RcEV-1084-7 | 17106 | 863582 | 2468.48 | 21612 | 21.49 | 7762.14 |
| RcEV-1084-8 | 16383 | 3196611 | 9540.48 | 26585 | 27.60 | 29511.04 |
| RcEV-1084-9 | 15480 | 4270119 | 13487.85 | 93519 | 102.76 | 42283.31 |
| RcEV-10125-1 | 17181 | 708947 | 2324.27 | 21298 | 28.20 | 6375.46 |
| RcEV-10125-2 | 17274 | 38209 | 124.59 | 4092 | 5.39 | 367.32 |
| RcDMV-0928-1 | 3946 | 28695 | 434.51 | 1179470 | 6660.57 | 45926.19 |
| RcDMV-0928-2 | 3872 | 381338 | 5884.73 | 8480873 | 48807.55 | 343319.12 |
| RcDMV-0928-3 | 3737 | 21852 | 349.40 | 1150309 | 6859.20 | 47049.55 |
| RcDMV-0928-4 | 3436 | 66593 | 1158.05 | 2443837 | 15848.96 | 109593.86 |
| RcDMV-0928-5 | 2809 | 23694 | 504.01 | 880022 | 6981.10 | 48258.24 |
| RcDMV-0928-6 | 2674 | 79007 | 1765.45 | 1097027 | 9141.93 | 65970.49 |
| RcDMV-0942-1 | 3871 | 26 | 0.41 | 372 | 2.35 | 15.42 |
| RcDMV-0942-2 | 3723 | 4 | 0.07 | 229 | 1.50 | 9.39 |
| RcDMV-0942-3 | 3273 | 66 | 1.23 | 197 | 1.47 | 12.05 |
| RcDMV-0942-4 | 3180 | 0 | 0.00 | 326 | 2.50 | 15.38 |
| RcDMV-0942-5 | 2824 | 27814 | 598.99 | 3993280 | 34511.81 | 213585.02 |
| RcMV1 | 3149 | 119841 | 1860.83 | 1453445 | 7850.73 | 74942.17 |
| RcDMV-1084-2 | 3787 | 10758 | 138.90 | 1704059 | 7653.73 | 67922.51 |
| RcDMV-1084-3 | 3948 | 13284 | 164.52 | 3189683 | 13742.14 | 121693.28 |
| RcDMV-1084-4 | 3630 | 93124 | 1254.38 | 3385805 | 15864.97 | 143757.40 |
| RcDMV-1084-5 | 3244 | 85329 | 1286.14 | 7023255 | 36824.89 | 328695.31 |
| RcDMV-1084-6 | 2830 | 36411 | 629.10 | 1131394 | 6800.04 | 61897.79 |
| RcDMV-10125-1 | 4004 | 52245 | 734.97 | 3196091 | 18155.67 | 121690.91 |
| RcDMV-10125-2 | 3789 | 16348 | 243.03 | 2460922 | 14772.72 | 98070.86 |
| RcDMV-10125-3 | 3347 | 152952 | 2574.07 | 6554586 | 44542.73 | 300606.72 |
| RcDMV-10125-4 | 2848 | 122386 | 2420.54 | 3928163 | 31371.61 | 213336.50 |
| RcBYV-0928-1 | 13512 | 86828 | 383.97 | 1917 | 3.16 | 985.18 |
| RcBYV-0928-2 | 13459 | 82234 | 365.08 | 2166 | 3.59 | 940.63 |
| RcBYV-0928-3 | 13450 | 75112 | 333.69 | 1884 | 3.12 | 858.69 |
| RcBYV-0928-4 | 13389 | 81287 | 362.76 | 2519 | 4.19 | 938.90 |
| RcBYV-0928-5 | 12874 | 13432 | 62.34 | 591 | 1.02 | 163.39 |
| RcBYV-0928-6 | 12868 | 11306 | 52.50 | 1026 | 1.78 | 143.75 |
| RcBYV-0928-7 | 12147 | 15718 | 77.32 | 1095 | 2.01 | 207.62 |
| RcBYV-0928-8 | 11774 | 53277 | 270.38 | 2062 | 3.90 | 705.02 |
| RcBYV-0942-1 | 13071 | 14420 | 67.09 | 3065 | 5.72 | 200.65 |
| RcBYV-0942-2 | 13038 | 29018 | 135.36 | 6815 | 12.76 | 412.25 |
| RcBYV-0942-3 | 12859 | 13527 | 63.98 | 2784 | 5.28 | 190.27 |
| RcBYV-1084-1 | 13394 | 13539 | 49.43 | 0 | 0.00 | 151.62 |
| RcBYV-1084-2 | 13000 | 7113 | 26.75 | 0 | 0.00 | 82.07 |
| RcBYV-1084-3 | 13026 | 80405 | 301.82 | 0 | 0.00 | 925.90 |
| RcBYV-1084-4 | 12981 | 68532 | 258.14 | 0 | 0.00 | 791.91 |
| RcBYV-1084-5 | 12912 | 7702 | 29.17 | 0 | 0.00 | 89.47 |
| RcBYV-10125-1 | 13531 | 44474 | 185.14 | 9883 | 16.61 | 602.58 |
| RcBYV-10125-2 | 13528 | 46508 | 193.65 | 13991 | 23.52 | 670.82 |
| RcBYV-10125-3 | 12997 | 10541 | 45.68 | 6648 | 11.63 | 198.38 |
| RcBYV-10125-4 | 12849 | 12473 | 54.68 | 10658 | 18.87 | 270.03 |
| RcLeV-0928-1 | 8771 | 15620 | 106.41 | 5792 | 14.72 | 366.18 |
| RcLeV-0928-2 | 7080 | 12047 | 101.67 | 3956 | 12.45 | 339.05 |
| RcLeV-0928-3 | 7016 | 9889 | 84.22 | 3000 | 9.53 | 275.56 |
| RcLeV-0928-4 | 7011 | 15339 | 130.73 | 3070 | 9.76 | 393.86 |
| RcPhV-0928-1 | 11394 | 64726 | 339.43 | 336 | 0.66 | 856.53 |
| RcPhV-0928-2 | 9856 | 34028 | 206.29 | 1826 | 4.13 | 545.67 |
| RcPhV-0928-3 | 6445 | 100264 | 929.55 | 1133 | 3.92 | 2359.90 |
| RcPhV-0942-1 | 9478 | 574 | 3.68 | 1256 | 3.23 | 28.96 |
| RcPhV-0942-2 | 6457 | 90590 | 853.24 | 3262 | 12.33 | 2180.24 |
| RcPhV-1084-1 | 10613 | 19449 | 89.61 | 56320 | 90.26 | 1070.89 |
| RcPhV-1084-2 | 9477 | 9385 | 48.42 | 3430 | 6.16 | 202.83 |
| RcPhV-1084-3 | 9378 | 38164 | 198.98 | 6556 | 11.89 | 715.29 |
| RcPhV-10125-1 | 11079 | 72318 | 367.68 | 17547 | 36.02 | 1216.69 |
| RcPhV-10125-2 | 10547 | 264842 | 1414.42 | 36398 | 78.49 | 4284.25 |
| RcPhV-10125-3 | 4624 | 81476 | 992.50 | 739 | 3.64 | 2667.01 |
| RcOCV-0928-1 | 2149<br>(RNA1) | 131729 | 3662.66 | 127 | 1.32 | 9203.54 |
|  | 1789<br>(RNA2) | 224587 | 7501.12 | 84 | 1.05 | 18837.70 |
| RcOCV-0928-2 | 2222<br>(RNA1) | 77729 | 2090.22 | 808 | 8.10 | 5301.78 |
|  | 1750<br>(RNA2) | 77049 | 2630.76 | 36 | 0.46 | 6607.29 |
| RcOCV-0928-3 | 2285<br>(RNA1) | 394717 | 10321.71 | 168 | 1.64 | 25922.43 |
|  | 1778<br>(RNA2) | 160181 | 5383.09 | 879 | 11.02 | 13587.74 |
| RcOCV-0928-4 | 2252<br>(RNA1) | 315499 | 8371.08 | 126 | 1.25 | 21022.98 |
|  | 1767<br>(RNA2) | 208255 | 7042.24 | 485 | 6.12 | 17719.86 |
| RcOCV-0942-1 | 2238<br>(RNA1) | 698913 | 18992.61 | 2741 | 29.89 | 47027.75 |
|  | 1773<br>(RNA2) | 208284 | 7144.45 | 1597 | 21.98 | 17756.43 |
| RcOCV-0942-2 | 2148<br>(RNA1) | 70366 | 1992.28 | 365 | 4.15 | 4939.32 |
|  | 1750<br>(RNA2) | 168 | 5.84 | 0 | 0.00 | 14.40 |
| RcOCV-0942-3 | 2192<br>(RNA1) | 209 | 5.80 | 0 | 0.00 | 14.30 |
|  | 1741<br>(RNA2) | 134428 | 4695.83 | 2194 | 30.76 | 11770.99 |
| RcOCV-1084-1 | 2230<br>(RNA1) | 191037 | 4188.77 | 208 | 1.59 | 12864.01 |
|  | 1872<br>(RNA2) | 56971 | 1488.06 | 216 | 1.96 | 4582.29 |
| RcOCV-1084-2 | 2225<br>(RNA1) | 406794 | 8939.61 | 405 | 3.10 | 27451.62 |
|  | 1815<br>(RNA2) | 128061 | 3449.96 | 611 | 5.73 | 10634.05 |
| RcOCV-1084-3 | 2154<br>(RNA1) | 64222 | 1457.85 | 4 | 0.03 | 4472.56 |
|  | 1722<br>(RNA2) | 354961 | 10079.09 | 181 | 1.79 | 30935.71 |
| RcOCV-1084-4 | 2015<br>(RNA1) | 482217 | 11701.49 | 120 | 1.01 | 35905.98 |
|  | 1661<br>(RNA2) | 475507 | 13997.84 | 348 | 3.56 | 42973.06 |
| RcOCV-10125-1 | 2239<br>(RNA1) | 511373 | 12864.83 | 83 | 0.84 | 34264.58 |
|  | 1774<br>(RNA2) | 392587 | 12465.30 | 54 | 0.69 | 33199.63 |
| RcOCV-10125-2 | 2238<br>(RNA1) | 129605 | 3261.99 | 294 | 2.99 | 8706.37 |
|  | 1749<br>(RNA2) | 12025 | 387.27 | 48 | 0.62 | 1035.42 |
| RcOCV-10125-3 | 2163<br>(RNA1) | 427463 | 11131.72 | 722 | 7.59 | 29693.83 |
|  | 1732<br>(RNA2) | 730147 | 23745.58 | 3353 | 44.03 | 63524.83 |
| RcHV-0928-1 | 20697 | 18623 | 53.76 | 14970 | 16.12 | 243.46 |
| RcHV-0942-1 | 20803 | 21482 | 62.80 | 48957 | 57.44 | 507.90 |
| RcHV-1084-1 | 20713 | 14648 | 34.58 | 21616 | 17.75 | 262.62 |
| RcZHV-0928-1 | 13262 | 332865 | 1499.72 | 133124 | 223.68 | 5270.57 |
| RcZHV-0928-2 | 13292 | 387096 | 1740.13 | 152218 | 255.19 | 6086.15 |
| RcZHV-0942-1 | 13274 | 528 | 2.42 | 80 | 0.15 | 6.87 |
| RcZHV-1084-1 | 13292 | 435766 | 1603.01 | 573747 | 734.20 | 11392.34 |
| RcZHV-10125-1 | 14723 | 463650 | 1773.84 | 134708 | 208.11 | 6096.16 |
| RcZHV-10125-2 | 12321 | 417752 | 1909.82 | 372283 | 687.25 | 9618.15 |
| RcZHV-10125-3 | 13337 | 564243 | 2383.02 | 1000790 | 1706.76 | 17601.78 |
| RcPuV-1084-1 | 6728 | 8924 | 64.86 | 0 | 0.00 | 198.96 |
| RcPuV-10125-1 | 6641 | 8217 | 69.69 | 258 | 0.88 | 191.42 |

### Novel dsRNA viruses in the phylum *Duplornaviricota*

Three megabirnavirus-like dsRNA viruses in the phylum *Duplornaviricota*, order *Ghabrivirales*, were identified in strain R0928. The phylogenetic tree was reconstructed on the basis of the amino acid sequence of RdRp regions (Fig. 1 A). Representative viruses in the *Quadriviridae, Totiviridae, Chrysoviridae* and *Megabirnaviridae* families in the order *Ghabrivirales* and some viruses selected based on BLAST searches were analyzed. Many unclassified viruses closely related to the *Megabirnaviridae* family clustered on at least five branches, most of which were mycoviruses. Three mycoviruses in strain R0928 and other viruses, including Ceratobasidium megabirnavirus-like, Rhizoctonia solani megabirnavirus 1 and 2, Pterostylis megabirnavirus-like and Ceratobasidium mycovirus-like virus, clustered on a single branch with a bootstrap support value of 92% (Fig. 1 A). We named these three novel mycoviruses Rhizoctonia cerealis megabirnavirus-like virus-0928–1, −2 and −3, abbreviated as RcMBLV-0928–1, −2 and −3.

**Figure 1.**
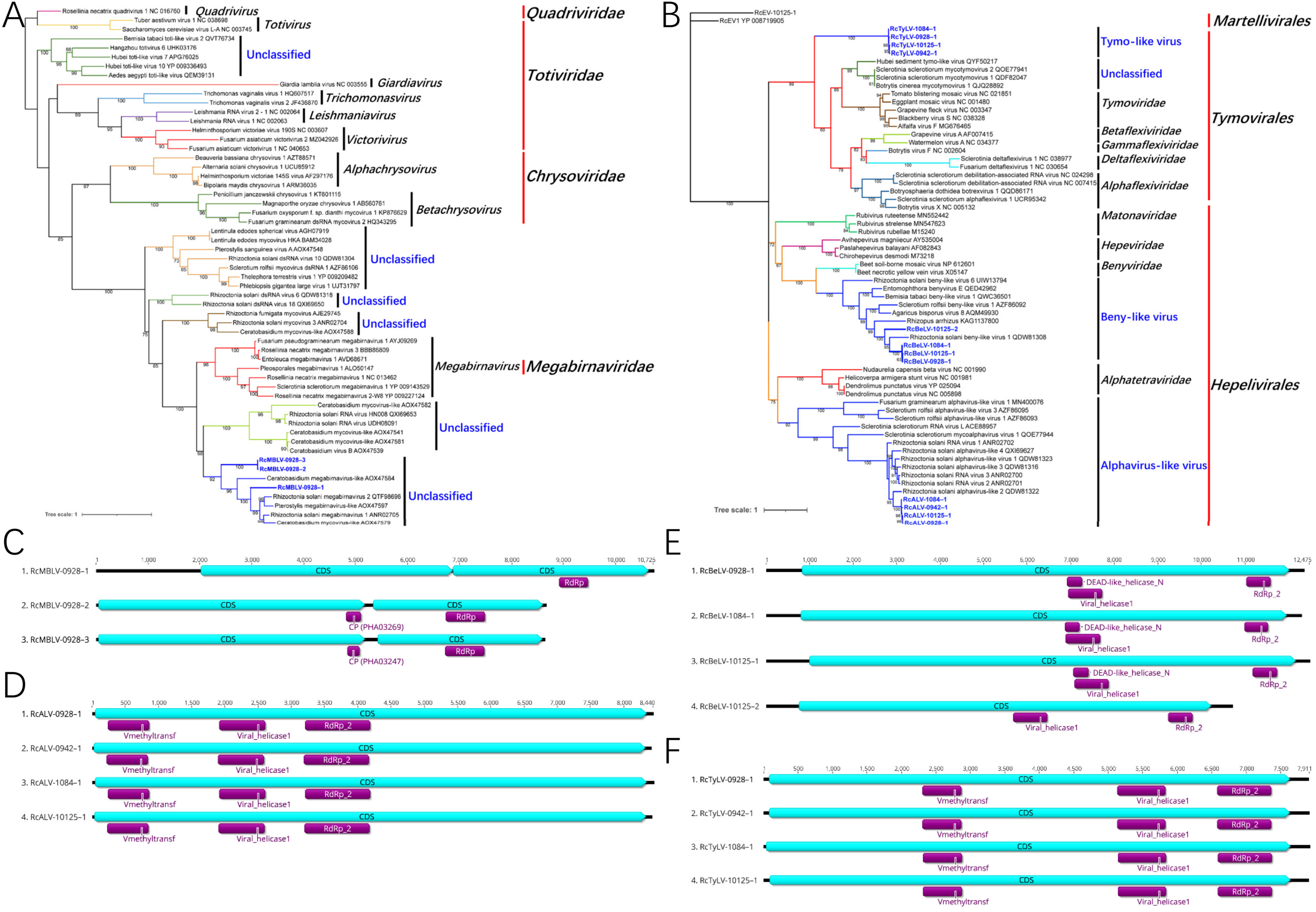
ML phylogenetic trees based on the RdRp regions and genome organization of the novel viruses identified from *R. cerealis* in the phyla *Duplornaviricota and Kitrinoviricota*. (A) Phylogenetic tree of megabirnavirus-like viruses constructed with the best-fit model “LG+R5+F”. (B) Phylogenetic tree of viruses in the phylum *Kitrinoviricota* constructed with the best-fit model “LG+I+G4+F”. Genome organization of megabirnavirus-like viruses (C), alphavirus-like viruses (D), beny-like virus (E) and tymo-like viruses (F).

All 3 viruses contained 2 ORFs that were predicted to encode a putative polyprotein. The genome lengths of these 3 viruses ranged from 8 626 bp to 10 725 bp, and all of them contained 2 coding sequences (CDSs) that were predicted to encode a putative polyprotein (Table 1; Fig. 1 C). Based on a CD search, the viral RdRp domain was located in the second CDS, and 2 viruses contained a coat protein (CP) domain in the first CDS (Fig. 1 C). The viruses in the family *Megabirnaviridae* had two genome segments. However, the genome of RcMBLVs in *R. cerealis* and some other megabirnavirus-like viruses in the same clade in the phylogenetic tree only had one segment (Fig. 1 A C). The pairwise identities of the amino acid sequences of the RdRp regions of RcMBLVs compared with the viruses in the *Megabirnaviridae* family were less than 40%, which indicated the identification of a new family in the order *Ghabrivirales*.

### Novel ssRNA(+) viruses in the phylum *Kitrinoviricota*

In the four *R. cerealis* strains, we identified 32 ssRNA(+) mycoviruses in the phylum *Kitrinoviricota*. Among these viruses, 17 alphaendornaviruses and 3 betaendornaviruses had been reported previously [22]. Based on the phylogenetic analysis, 8 novel viruses belonged to the *Hepelivirales* order (Fig. 1 B), 4 of which were named Rhizoctonia cerealis alphavirus-like virus-0928–1, −0942–1, −1084–1 and −10125– 1, which were abbreviated as RcALV-0928–1, −0942–1, −1084–1 and −10125–1. The genome lengths of these 4 viruses ranged from 8 402 bp to 8 440 bp. All 4 RcALVs contained one CDS that was predicted to encode a putative polyprotein, and Vmethyltransf, viral helicase1 and RdRp_2 domains were located in this CDS (Fig. 1 D). Four novel viruses were named Rhizoctonia cerealis beny-like virus-0928–1, −0942–1, - 1084–1 and −10125–1, which were abbreviated as RcBeLV-0928–1, −0942–1, −1084–1 and −10125–1. The genome lengths of these 4 viruses ranged from 10 701 bp to 12 475 bp. These 4 RcBeLVs contained one CDS, and viral helicase1 and RdRp_2 domains were located in this CDS (Fig. 1 E).

In the phylogenetic tree, the 4 RcALVs and other mycoviruses from *R. solani, Sclerotinia sclerotiorum, S. rolfsii* and *Fusarium graminearum* clustered on a branch with a bootstrap support value of 92%. This Alphavirus-like virus group was the sister group to the *Alphatetraviridae* family in the *Hepelivirales* order (Fig. 1 B). The 4 RcBeLVs and other mycoviruses clustered on a branch with a bootstrap support value of 100%, which formed the Beny-like virus group. The phylogenetic position of this group was closely related to the *Benyviridae* family (Fig. 1 B). In the order *Hepelivirales*, the alphavirus-like and beny-like virus groups represented two novel groups at the family level.

The other 4 novel viruses were identified as tymo-like viruses in the order *Tymovirales* (Fig. 1 B). We named them Rhizoctonia cerealis tymo-like virus-0928–1, −0942–1, −1084–1 and −10125–1, which were abbreviated as RcTyLV-0928–1, −0942–1, - 1084–1 and −10125–1. The genome lengths of these 4 viruses ranged from 7 888 bp to 7 911 bp. All 4 RcTyLVs contained one CDS, and Vmethyltransf, viral helicase1 and RdRp_2 domains were located in this CDS (Fig. 1 F). The 4 RcTyLVs from *R. cerealis* were closely related to each other, and their genomic identities ranged from 79.8-81.8%. These 4 viruses clustered on a separate branch in the phylogenetic tree (Fig. 1 B). Using the amino acid sequences of the RdRp region as queries in BLASTp analysis, the closest viruses identified were Sclerotinia sclerotiorum mycotymovirus 1 and 2 and Botrytis cinerea mycotymovirus 1, which belonged to the unclassified group in the *Tymovirales* order. Based on phylogenetic analysis, the tymo-like virus group was independent of the *Alphaflexiviridae*, *Betaflexiviridae*, *Gammaflexiviridae*, *Deltaflexiviridae* and *Tymoviridae* families, which indicates the identification of a new family in the *Tymovirales* Order.

### Novel ssRNA(+) viruses in the phylum *Lenarviricota*

Twenty-one mitoviruses with ssRNA(+) genomes were identified in the four *R. cerealis* strains. Based on the phylogenetic analysis, all of these mitoviruses were identified as *Duamitovirus* in the family *Mitoviridae*, order *Cryppavirales*, phylum *Lenarviricota* (Fig. 2 A). We named them Rhizoctonia cerealis duamitovirus, abbreviated as RcDMV. RcMV1 in strain R1084 has been reported previously [21], and we retained the name RcDMV-1084–1 for this virus (Table 1). The genome lengths of these 21 mitoviruses ranged from 2 674 bp to 4 004 bp. These RcDMVs contained one CDS, and Mitovirus RNA-dependent RNA polymerase domains were located in this CDS (Fig. 2 B). However, in the phylogenetic tree, the mitoviruses from *R. cerealis* did not cluster together but were dispersed in multiple clades, indicating that these RcDMVs showed rich diversity at the species level.

**Figure 2.**
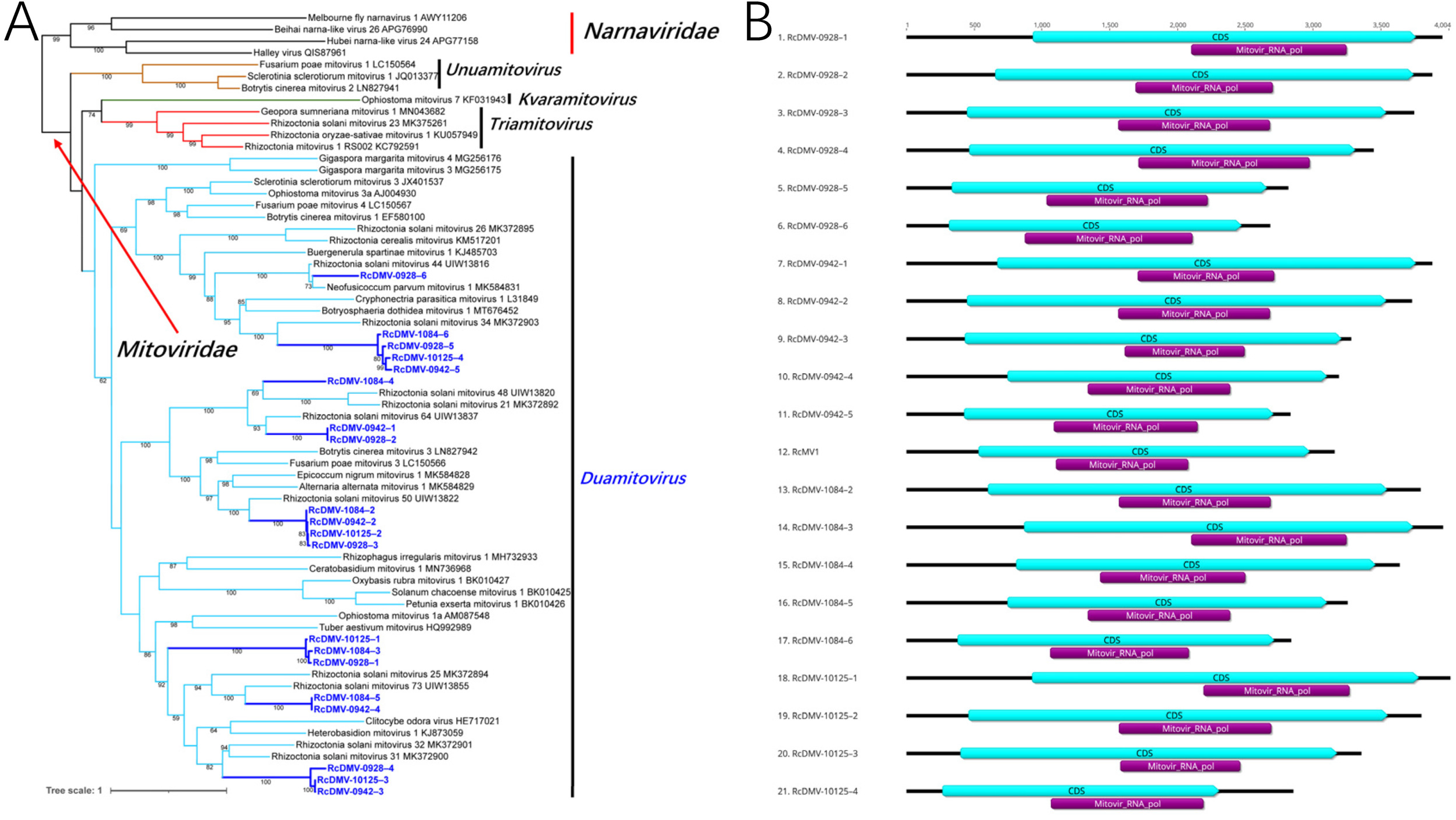
ML phylogenetic trees based on the RdRp regions and genome organization of the novel viruses identified from *R. cerealis* in the family *Mitoviridae*. (A) Phylogenetic tree of mitoviruses constructed with the best-fit model “Blosum62+R6+F”. (B) Genome organization of mitoviruses.

### Novel ssRNA(-) viruses in the phylum *Negarnaviricota*

In total, 35 novel ssRNA(-) viruses in the *Negarnaviricota* phylum were identified in the four *R. cerealis* strains (Table 1, Fig. 3 A B C). Twenty viruses were identified as novel members of the *Bunyavirales* order (Fig. 3 A). These viruses were named Rhizoctonia cerealis bunyavirus, abbreviated as RcBYV. The genome lengths of these 20 RcBYVs ranged from 11 774 bp to 13 531 bp. With the exception of RcBYV-0928–7, the other 19 viruses contained 2 ORFs encoding two CDSs, and the Bunyavirus RNA-dependent RNA polymerase (Bunya_RdRp) domain was located in the first CDS (Fig. 3 D). Some of the viruses also contained a PIN_SF domain, which belongs to a large nuclease superfamily, in the first CDS (Fig. 3 D). However, no conserved domains have been found in the second CDS thus far. In the phylogenetic tree, all of the RcBYVs and Rhizoctonia solani Khurdun virus were clustered in a clade with a bootstrap support value of 99% (Fig. 3 A). This group was independent of other families in the *Bunyavirales* order and formed sister groups with the family *Sclerobunyaviridae*, which was proposed in 2021 [33]. At present, all of the viruses in this group were identified in *Rhizoctonia* fungi, so we proposed a new family *Rhizoctobunyaviridae* and a new genus *Rhizoctobunyavirus* to accommodate these viruses.

**Figure 3.**
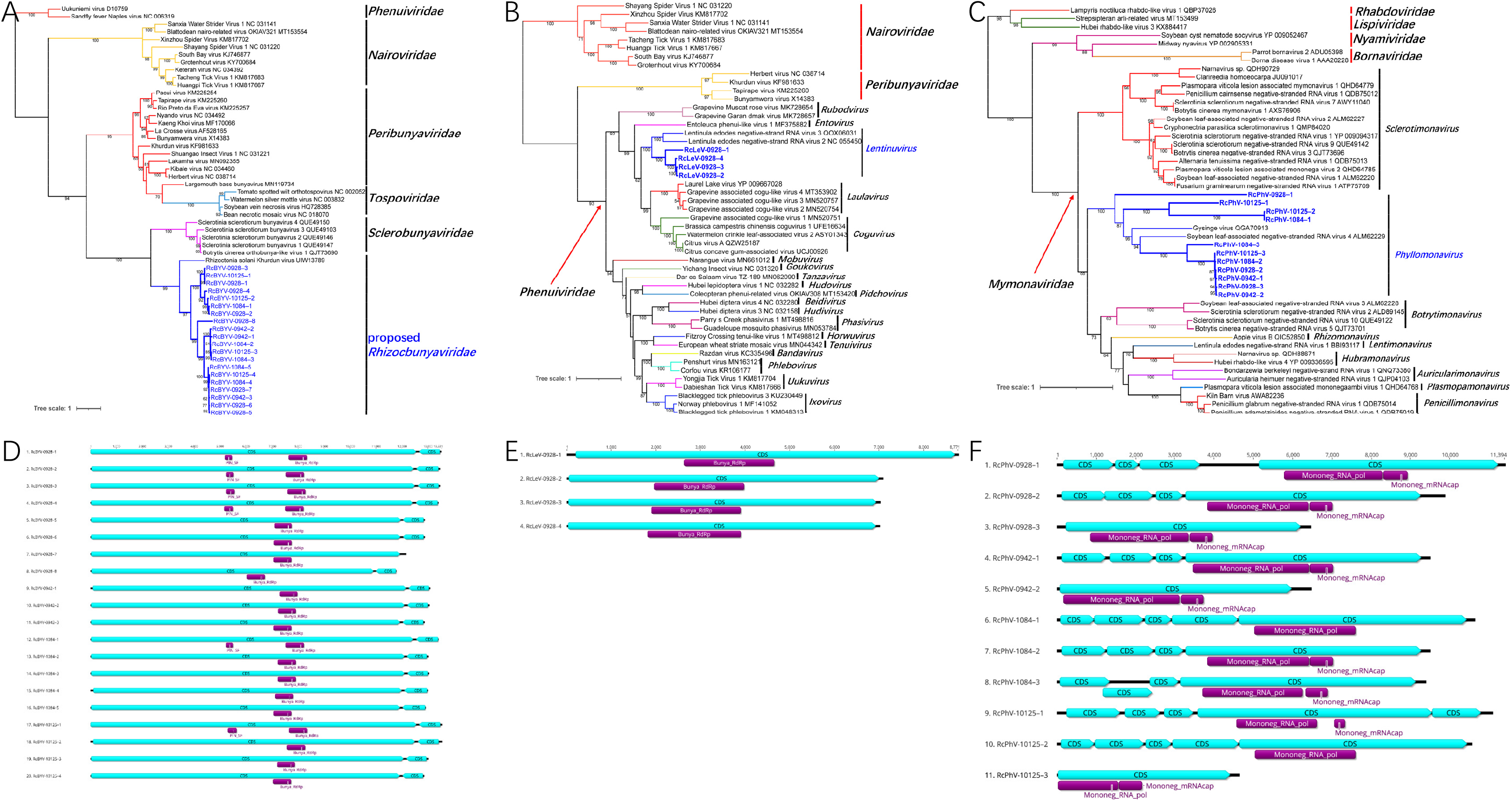
ML phylogenetic trees based on the RdRp regions and genome organization of the novel viruses identified from *R. cerealis* in the phylum *Negarnaviricota*. (A) Phylogenetic tree of bunyaviruses constructed with the best-fit model “LG+I+G4+F”. (B) Phylogenetic tree of lentinuviruses constructed with the best-fit model “LG+I+G4+F”. (C) Phylogenetic tree of phyllomonaviruses constructed with the best-fit model “LG+R5”. Genome organization of bunyaviruses (D), lentinuviruses (E) and phyllomonaviruses (F).

**Figure 4.**
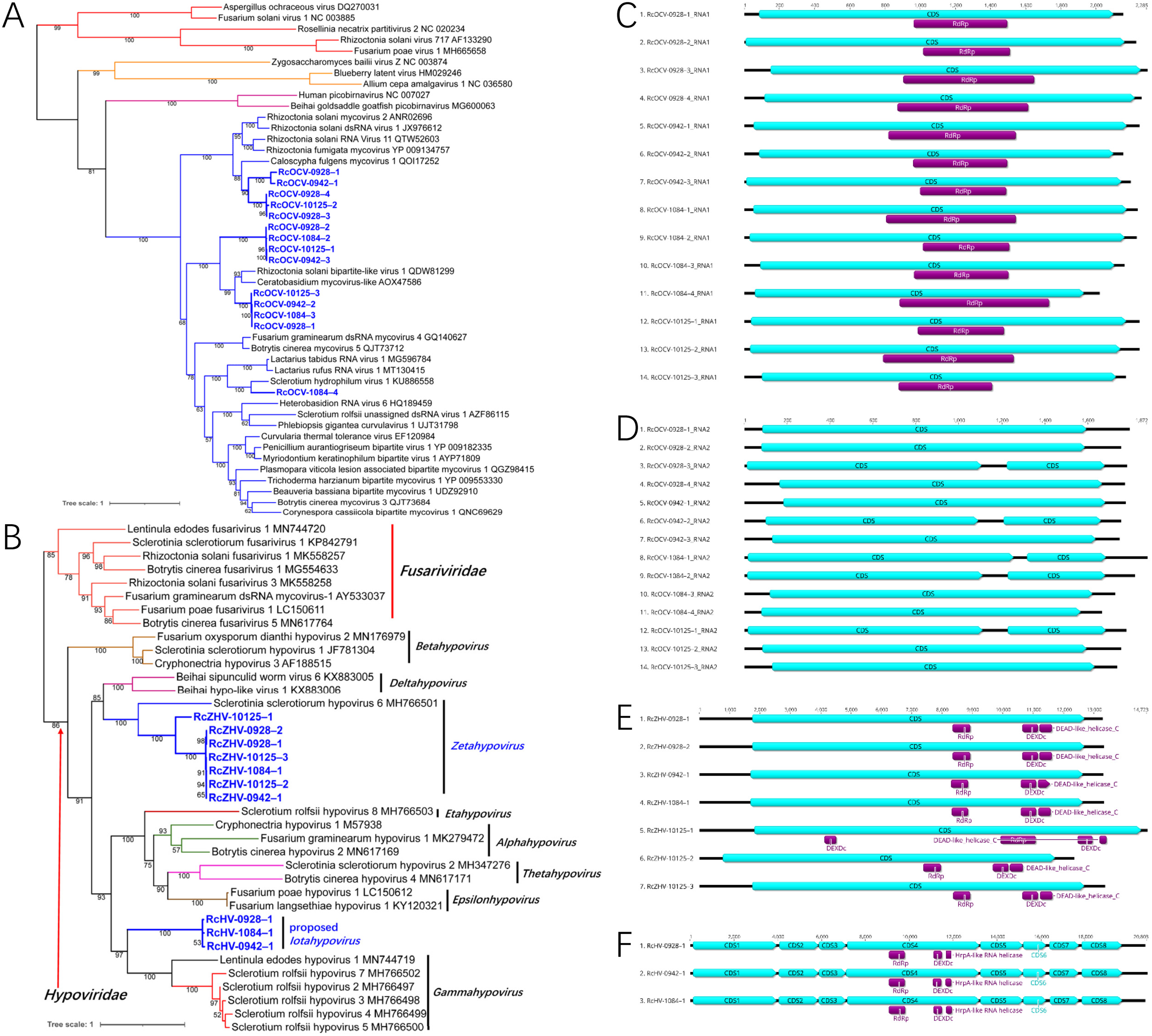
ML phylogenetic trees based on the RdRp regions and genome organization of the novel viruses identified from *R. cerealis* in the phylum *Durnavirales*. (A) Phylogenetic tree of orthocurvulaviruses constructed with the best-fit model “LG+I+G4+F”. (B) Phylogenetic tree of hypoviruses constructed with the best-fit model “Blosum62+R3”. (C) Genome organization of orthocurvulavirus RNA1 (C) and RNA2 (D), *Zetahypovirus* (E) and a proposed *Iotahypovirus* (F).

Four ssRNA(-) viruses in strain R0928 were identified as lentinuviruses in the *Phenuiviridae* family in the *Bunyavirales* order (Fig. 3 B). These viruses were named Rhizoctonia cerealis lentinuvirus, abbreviated as RcLeV. The genome lengths of these 4 RcLeVs ranged from 7 011 bp to 8 771 bp. All of the RcLeVs contained one CDS, and the Bunya_RdRp domain was located in this CDS (Fig. 3 E). Based on the phylogenetic analysis, the 4 RcLeVs clustered into a single clade and then clustered into a larger clade with Lentinula edodes negative-strand RNA 2 and 3 (LeNSRV2 and LeNSRV3) in the *Lentinuvirus* genus (Fig. 3 B). This larger clade was independent of other genera in the *Phenuiviridae* family with 100% bootstrap support (Fig. 3 B). Therefore, we classified these 4 viruses from *R. cerealis* as members of the *Lentinuvirus* genus. However, the genetic identity of the RdRp regions among the 4 RcLeVs and 2 LeNSRVs was less than 60%, so the possibility of establishing a new genus in the future could not be ruled.

The other 11 novel ssRNA(-) viruses were identified as new *Phyllomonavirus* members belonging to the *Mymonaviridae* family in the *Mononegavirales* order (Fig. 3 C). These viruses were named Rhizoctonia cerealis phyllomonavirus, abbreviated as RcPhV. The genome structures of these viruses were diverse, and most of them contained 4 or 5 CDSs. However, RcPhV-0928–3, RcPhV-0942–2 and RcPhV-10125–3, with shorter genome lengths (6 445, 6 457 and 4 624 bp, respectively), only contained one CDS (Fig. 3 F). The genome lengths of other RcPhV viruses ranged from 9 378 bp to 11 394 bp. All of the RcPhVs contained a Mononeg_RNA_pol domain located in the longest CDS, and some of them also had a Mononeg_mRNAcap domain located after the Mononeg_RNA_pol domain (Fig. 3 F). In the phylogenetic tree, all of the RcPhVs, a Gysinge virus, and Soybean leaf-associated negative-stranded RNA virus 4 were clustered in a clade with bootstrap support value 100% (Fig. 3 C). The viruses in this clade belonged to the genus *Phyllomonavirus*, and this clade was independent of other genera in the *Mymonaviridae* family (Fig. 6 B). In the *Phyllomonavirus* clade, the RcPhVs identified in *R. cerealis* were not tightly clustered onto one branch, indicating their diversity.

**Figure 5.**
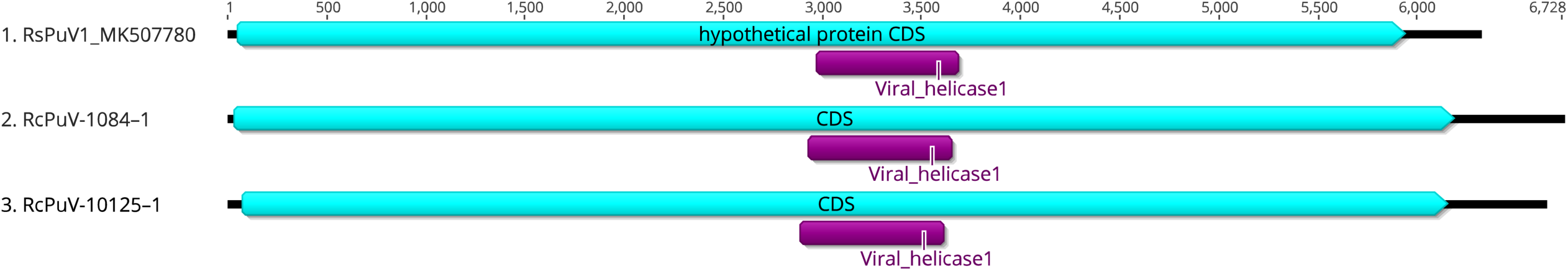
Genome organization of putative viruses identified in *R. cerealis* and RsPuV1 in *R. solani*.

**Figure 6.**
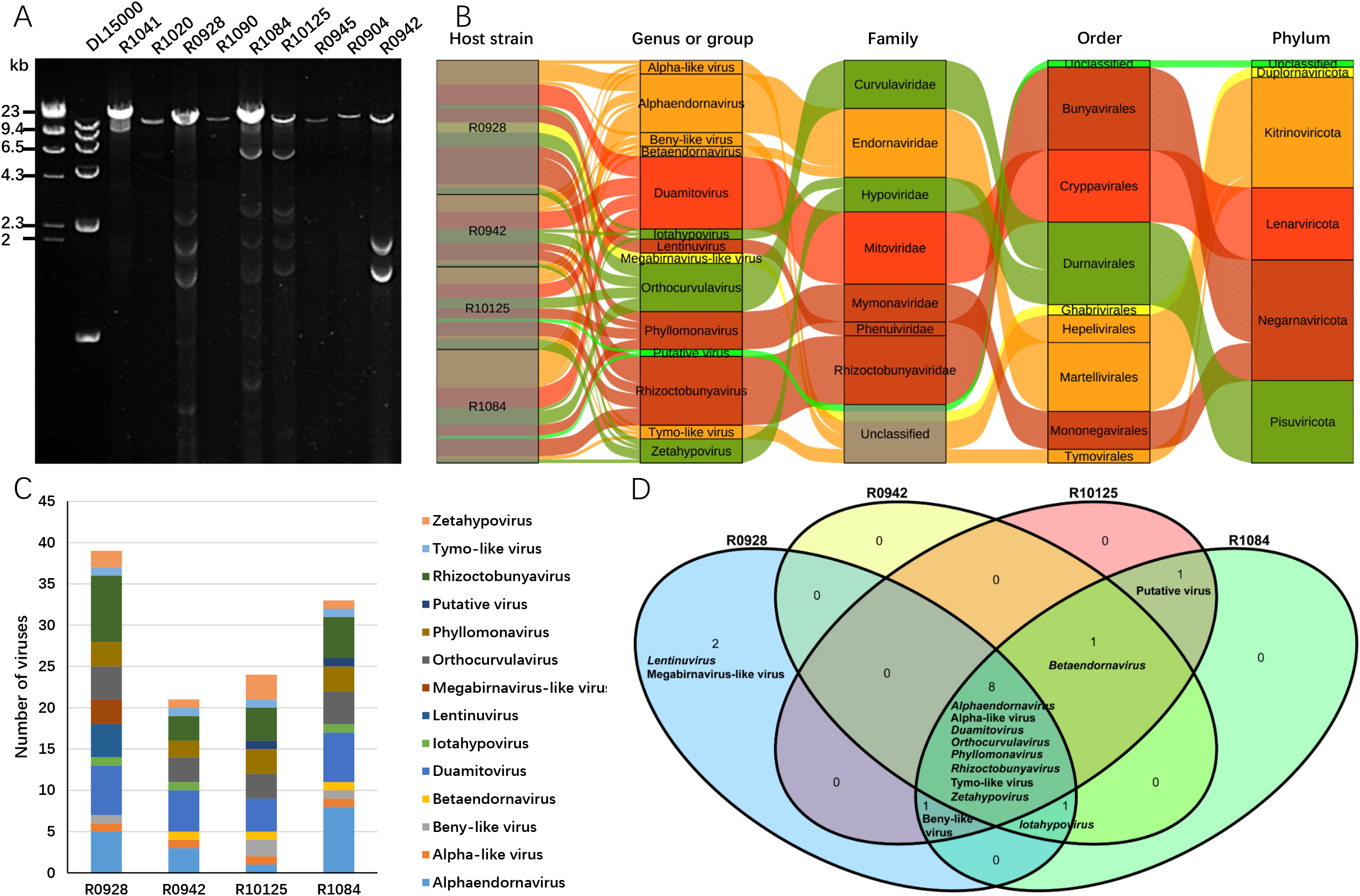
Diversity of viruses identified in *R. cerealis*. (A) Agarose gel electrophoresis of dsRNA extracted from different strains. Bands of the DL15000 marker: 15, 10, 7, 5, 2.5, and 1 kbp. (B) Sankey diagram showing the compositions of viruses from four strains. (C) Viruses belonging to different genera or groups presented in each strain. (D) The numbers and groups of shared viruses in the virome of four strains of *R. cerealis*.

### Novel dsRNA and ssRNA(+) viruses in the phylum *Pisuviricota*

Twenty-four novel dsRNA and ssRNA(+) viruses in the four *R. cerealis* strains were identified as belonging to the *Pisuviricota* phylum (Table 1, Fig. 4 A B). Among these viruses, 14 dsRNA viruses with 2 genome segments were identified as orthocurvulaviruses in the *Curvulaviridae* family, *Durnavirales* order (Fig. 4 A). These viruses were named Rhizoctonia cerealis orthocurvulavirus, abbreviated as RcOCV. The lengths of their RNA1 segments ranged from 2 015 bp to 2 285 bp, and the lengths of the RNA2 segments ranged from 1 661 bp to 1 778 bp. All of the RNA1 segments contained only one CDS, and the RdRp domain was located in this CDS (Fig. 4 C). Among the 14 RNA2 segments, 5 contained two CDSs, and the others contained only one CDS. There was no identifiable conserved domain within the RNA2 segment (Fig. 4 D). However, since there multiple RcOCV genomes could exist in the same strain, we could not match the RNA1 and RNA2 segments of each virus, so we temporarily named them according to the order in which they were discovered (Table 1, Fig. 4 C D).

Currently, there is only one genus, *Orthocurvulavirus*, in the family *Curvulaviridae*. Based on the topological structure of the phylogenetic tree, the viruses in the genus *Orthocurvulavirus* could be divided into 3 groups (Fig. 4 A). The hosts of viruses in groups I and II were mainly *Rhizoctonia* fungi, while the hosts of viruses in group III were various fungi. Among the 14 RcOCVs, 5 clustered in group I, 8 clustered in group II, and only RcOCV-1084–4 clustered in group III (Fig. 4 A). The results of this phylogenetic analysis also provide evidence for establishing more genera in the *Curvulaviridae* family.

Based on the phylogenetic analysis, 10 ssRNA(+) viruses were related to hypoviruses (Table 1, Fig. 4 B). Seven of them were identified as zetahypoviruses and were named Rhizoctonia cerealis zetahypovirus, abbreviated as RcZHV. The genome lengths of these 7 RcZHVs ranged from 12 321 bp to 14 723 bp. All of the RcZHVs contained one CDS, and RdRp, DEADc and DEAD-like_helicase_C domains were located in this CDS (Fig. 4 E). Three other unclassified hypovirus viruses were identified from strains R0928, R0942 and R1084 and were named Rhizoctonia cerealis hypovirus, abberviated as RcHV. The genome lengths of these 3 RcHVs were 20 697 bp, 20 803 bp and 20 713 bp, respectively, and they contained 8 CDSs. RdRp, DEXDc and HrpA-like RNA helicase domains were located in CDS4, which was the longest CDS in the genome (Fig. 4 F).

At present, there are 8 genera in the family *Hypoviridae*: *Alpha-, Beta-* to *Thetahypoviruses* [34]. In the phylogenetic tree, 7 RcZHVs and Sclerotinia sclerotiorum hypovirus 6 clustered in the *Zetahypovirus* clade with 100% bootstrap support (Fig. 4 B). The 3 RcHVs in *R. cerealis* were closely related to each other, and their genomic identities ranged from 72.5-76.8%. The RcHVs clustered in a separate clade, which was independent of other genera in the *Hypoviridae* family (Fig. 4 B). Based on their genome structure, which differed from those of other hypoviruses, and the results of phylogenetic analysis, we proposed a new genus, *Iotahypovirus*, to accommodate these 3 new hypoviruses from *R. cerealis*.

### Unclassified viral genome sequences

In strains R1084 and R10125, we identified two viral genome sequences that could not be assigned to any existing clade of viruses present in the databases. The lengths of these two sequences were 6 728 bp and 6 641 bp, and each contained one CDS. We could not identify an RdRp domain in the CDS, but a viral helicase 1 domain was located within it (Fig. 5). The pairwise identity between these two sequences was 98.1%. In BLAST searches using the genome sequences or the amino acid sequences of the Viral_helicase1 domain, the first match was Rhizoctonia solani putative virus 1 (RsPuV1), which was also identified in *Rhizoctonia* fungi. The genome structures were similar among these 3 viral genome sequences (Fig. 5); however, the pairwise identity of the Viral_helicase1 region between the viruses from *R. cerealis* and *R. solani* was less than 28.5%. We named these two viruses Rhizoctonia cerealis putative virus, which were abbreviated as RcPuV-1084–1 and RcPuV-10125–1.

### Diversity of mycoviruses present in the four strains of *R. cerealis*

The diversity of dsRNA bands in the R0928, R0942, R1084 and R10125 strains was particularly rich (Fig. 6 A). In these four strains, a total of 117 mycoviruses were found. These viruses belonged to at least 9 genera and 5 unclassified genera, 7 families and 3 unclassified families, and 5 phyla in the *Orthornavirae* kingdom (Fig. 6 B). At the phylum level, ssRNA(-) viruses in the *Negarnaviricota* phylum were the most abundant, accounting for 30% of the total. With the exception of unclassified viruses, the fewest identified dsRNA viruses belonged to the *Duplornaviricota* phylum, representing only 3% of the total. At the family level, viruses of the *Mitoviridae* and *Endornaviridae* families were the most abundant, followed by the proposed new family *Rhizoctobunyaviridae* (Fig. 6 B). At the genus level, the four strains contained a wide variety of viruses, among which R0928 and R1084 each contained 12 genera, R10125 contained 11 genera, and R0942 contained 10 genera. The R0928 strain harbored the greatest number of viruses, with 39 viruses found in this strain, while R0942 contained the fewest viruses, with 21 viruses found in this strain (Fig. 6 C). Viruses of 8 genera could be identified in all four strains, indicating that these viruses coinfected *Rhizoctonia* fungi. Betaendornaviruses, beny-like viruses, and iotahypoviruses were identified in 3 strains, putative viruses (RcPuV) were identified in R1084 and R10125, and lentinuviruses and megabirnavirus-like viruses were identified only in R0928 (Fig. 6 D).

## Discussion

At present, high-throughput sequencing has become the main method for discovering new viruses from a large number of samples [3–4, 33]. *Rhizoctonia* are soilborne fungi that mostly lack sexual reproduction, and many of these fungi are plant pathogens [35]. In this study, we used two methods to construct cDNA sequencing libraries and combined the two resulting high-throughput sequencing datasets to assemble relatively complete viral genomes of 117 viruses from four *R. cerealis* strains (Table 1). The depth of coverage values of these viral genome sequences were generally more than 100X (Table 2). Using the RPKM value of each viral genome sequence as a measure, we compared the quality of the data obtained via the two methods. For most viruses other than mitoviruses, the dsRNA-Seq method produced higher RPKM values. Compared with the rRNA-depleted RNA-Seq method, more reads from the viral genome were obtained using the dsRNA-Seq method because a large number of nonviral RNAs were removed in the pretreatment stage. For some viruses, such as RcMBLV-0928–3, RcALV-1084–1, and RcEV-1084–1, no reads could be obtained using the rRNA-depleted RNA-Seq method but complete genome sequences could be obtained using the dsRNA-Seq method (Table 2). Therefore, we recommend the use of purified dsRNA for building a high-throughput sequencing library to obtain viral genome sequences from fungal hyphae. Previously, we reported the identification of endornaviruses with extremely long genomes and 2 ORFs in the 4 investigated *R. cerealis* strains [22]. The results of this study further demonstrate the reliability of these viral genomes and indicate that only the dsRNA-Seq method can produce such complete ultralong viral genomes (Table 2).

Mitoviruses in the family *Mitoviridae* are ubiquitously identified in filamentous fungi that have ssRNA(+) genomes of 2.3 to 3.6 kb encoding only RdRps [36]. Capsidless mitoviruses localize to fungal mitochondria, within which their replication cycle is completed, consistent with the mitochondrial codon usage of fungi [37]. In this study, with the exception of RcDMV-0942–1 to −0942–4, for which relatively low read numbers were obtained by both methods, unusually high RPKM values were obtained for some other mitoviruses using the RNA-depleted RNA-Seq method, although higher RPKM values were also obtained for these viruses using the dsRNA-Seq method (Table 2). DsRNA is an intermediate in viral replication, and the genome of mitoviruses consists of ssRNA. The results of this work indicated that in host fungi, the density of mitoviruses was significantly higher than that of other viruses (Table 2). We speculate that mitochondria act as a natural protective harbor, helping mitoviruses avoid interference by cytoplasmic RNA silencing.

Twenty-one duamitoviruses were discovered, and each strain contained 4 to 6 mitoviruses (Table 1), indicating that in *R. cerealis*, both the variety and density of mitoviruses were very high. To the best of our knowledge, although many mitoviruses have been identified in fungi, investigations of the density of these viruses in host cells are rare. At present, only a few mitoviruses, such as Botrytis mitovirus 1 (BcMV1) and Sclerotinia sclerotiorum mitovirus 1 (SsMV1), are known to cause severe symptoms, including hypovirulence, in host fungi [38–39]. Therefore, many mitoviruses exist in *R. cerealis* mitochondria, and their biological functions are of considerable interest. However, knowledge of this topic is currently considerably lacking.

The 117 viruses with dsRNA, ssRNA(+) and ssRNA(-) genomes identified in *R. cerealis* could be grouped into the *Duplornaviricota, Kitrinoviricota, Lenarviricota, Negarnaviricota* and *Pisuviricota* phyla in the *Riboviria* realm (Table 1). Most of these viruses were sufficiently distant from those deposited in the databases to suggest that they qualify as possible new viral species, genera or even families. We proposed names for these novel mycoviruses based on the name of the host strain and the number of the virus identified in the strain. This method generates names includes the host strain name to avoid the infinite expansion of Nos. or the application of the same No. to similar viruses by different researchers.

*Bunyavirales* is one of the largest orders of negative-strand RNA viruses in the *Negarnaviricota* phylum, containing 14 families based on the ICTV report from 2021 (https://talk.ictvonline.org/). The hosts of most bunyaviruses are vertebrates, arthropods, and plants [40]. However, some newly identified bunyaviruses were discovered in the pathogenic fungus *S. sclerotiorum*, and two new families, *Mycophenuiviridae* and *Sclerobunyaviridae*, were proposed in 2021 [33]. In *R. cerealis*, we identified 24 novel mycoviruses whose genomes contain a conserved Bunya_RdRp domain (Fig. 3 A B). However, the Bunya_RdRp regions of these bunyaviruses in *Rhizoctonia* fungi shared less than 26% identity with those of *Sclerobunyaviridae*, indicating that they represent a new virus lineage. Therefore, we propose a new family, *Rhizoctobunyaviridae*, and a new genus, *Rhizoctobunyavirus*, to accommodate these bunyaviruses in *Rhizoctonia* fungi. Furthermore, some additional clades with high bootstrap support also existed, indicating that more than one genus may be contained in this family (Fig. 3 A). Genomes of bunyaviruses are typically tripartite, consisting of large (L), medium (M), and small (S) segments [41]. However, the genomes of bunyaviruses found in *Sclerotinia* [33] and *Rhizoctonia* fungi only contained one segment, which was significantly distant from the sequences of other bunyaviruses (Fig. 3 D E).

The *Mymonaviridae* family includes many mycoviruses belonging to the genera *Sclerotimonavirus, Botrytimonavirus* and *Penicillimonavirus* [42]. In this study, 11 RcPhVs were discovered in *R. cerealis*, which were identified as phyllomonaviruses in the *Mymonaviridae* family (Fig. 3 C). This is the first report of mycoviruses in the genus *Phyllomonavirus*. The genome structures of the mycoviruses in the *Mymonaviridae* family were diverse. For example, the genome of Sclerotinia sclerotiorum negative-stranded RNA virus 1 strain AH98 contained 6 CDSs [43], and that of Botrytis cinerea mymonavirus 1 isolate Ecan17-2 contained 3 CDSs [44]. Similarly, 8 RcPhVs contained 4 to 5 CDSs in their genomes, and the other 3 RcPhV genomes contained only 1 CDS (Fig. 3 F).

In *R. solani*, Picarelli et al. [20] reported 4 viral fragments designated RsPuV 1 to 4, and the genome structures were similar to those of RcPuV-1084–1 and RcPuV-10125–1 from *R. cerealis*. We confirmed the presence of these two sequences based on the depth of coverage (greater than 190) and RT–PCR analysis (Table 2). These putative viruses only contained a Viral_helicase1 domain. Recent studies have indicated that Viral_helicase1 can undergo horizontal gene transfer between different viruses and/or bacteria [45]. At present, we cannot find any more information about these two sequences. They may be part of the genome of a virus for which the complete genome has not yet been obtained, or they may be fragments of multipartite virus genomes.

*R. cerealis* has no sexual reproductive stage and does not produce any asexual spores [35]. This means that the viruses contained in *R. cerealis* can only be transmitted by hyphal fusion or hyphal subgeneration. However, it is not clear how often hyphal fusion occurs in nature. Therefore, the transmission pattern of the viruses between different *R. cerealis* strains in nature is not clear. The analysis of dsRNA gel electrophoresis bands in different strains (Fig. 6 A) indicated that not all strains contained such rich diversity of mycoviruses, indicating that the transmission of viruses between different strains was not unimpeded. The four strains of *R. cerealis* investigated in this study were collected from three provinces in China with distant geographical locations. The virus species contained in these strains were different, and most of the genome sequences found even within the same species were different. These results also indicated that the evolution of viruses in different strains was relatively independent.

To our knowledge, many fungi harbor rich mycovirus diversity [20, 33], but there are few examples of fungal strains containing such rich diversity and large numbers of viruses. Based on RPKM values, we estimated the density of myoviruses in fungal host cells, for the first time. The results indicated that the density of some mycoviruses, such as mitoviruses, in host cells was very high (Table 2). Moreover, the evolutionary relationships among the many viruses coexisting within a single strain are of great interest. Highly similar sequences were rare among different viruses in the same strain, indicating that horizontal transfer or recombination of genome segments between different viruses was rare. At present, it is clear that mitoviruses exist only in fungal mitochondria [37], but the ecological niches of other viruses in host cells are not clear. Whether the different niche locations of viruses in fungal cells can reduce the occurrence of horizontal gene transfer among viruses needs further study.

In conclusion, the diversity of mycoviruses identified in *R. cerealis* was extremely rich, and many of these viruses were previously unreported. We demonstrated that a large number of mycoviruses can coexist in the same *R. cerealis* strain. This study provides a valuable resource for the subsequent utilization of mycoviruses and raises many new questions. We currently know very little about how these mycoviruses coexist in the same fungal cell, how they escape cytoplasmic RNA silencing by the host, and the occurrence of horizontal gene transfer, recombination and coevolution between different viruses.

## Acknowledgements

This work was supported by the National Natural Science Foundation of China (Grant No. 32072375) and the earmarked fund of the China Agriculture Research System (CARS-3) The authors have no conflicts of interest to declare.

